# Cis-Regulatory Atlas in Primary Human CD4+ T Cells

**DOI:** 10.1101/2022.12.09.519788

**Authors:** Kurtis Stefan, Artem Barski

**Affiliations:** Division of Allergy & Immunology, Cincinnati Children’s Hospital Medical Center, Cincinnati, OH 45229-3026, USA; Medical Scientist Training Program (MSTP), University of Cincinnati College of Medicine, Cincinnati, OH 45267, USA; Division of Human Genetics, Cincinnati Children’s Hospital Medical Center, Cincinnati, OH 45229-3026, USA; Department of Pediatrics, University of Cincinnati College of Medicine, Cincinnati, OH 45267, USA

**Keywords:** Cis-Regulatory Elements, Enhancers, Negative Regulatory Elements, Silencers, CD4, STARR-Seq

## Abstract

Cis-regulatory elements (CRE) are critical for coordinating gene expression programs that dictate cell-specific differentiation and homeostasis. Recently developed self-transcribing active regulatory region sequencing (STARR-Seq) has allowed for genome-wide annotation of functional CREs. Despite this, STARR-Seq assays are only employed in cell lines, in part, due to difficulties in delivering reporter constructs. Herein, we implemented and validated a STARR-Seq–based screen in human CD4+ T cells using a non-integrating lentiviral transduction system. Lenti-STARR-Seq is the first example of a genome-wide assay of CRE function in human primary cells, identifying thousands of functional enhancers and negative regulatory elements (NREs) in human CD4+ T cells. Results of the screen were validated using traditional luciferase assays. Genome-wide, we find clear differences between enhancers and NREs in nucleosome positioning, chromatin modification, eRNA production, and transcription factor binding. Our findings support the idea of silencer repurposing as enhancers in alternate cell types. Collectively, these data suggest that Lenti-STARR-Seq is a can be used for CRE screening in primary human cell types.

## Introduction

Cis-regulatory elements (CRE) are DNA elements that manage or modulate gene transcription. Putative CREs are frequently identified in structural assays, such as those measuring chromatin accessibility (e.g., DNase sequencing [DNase-Seq], assay for transposase-accessible chromatin sequencing [ATAC-Seq]), but these assays do not provide information about CRE function. CREs contribute to defining cell-specific gene expression programs through positive and negative regulation, and by insulating genes from inappropriate regulation. Annotation of CRE function is an ongoing challenge using existing datasets; the chromatin and genomic landscape of CREs is highly varied.(4, 5) Multiple attempts to identify CREs from existing data rely on correlating epigenetic chromatin opening and histone enrichment with function.(6–9) structure-based approach fails to explain expression changes: in our recent study of T cell activation, genes with nearby chromatin opening were found to be both up- and down-regulated.(10) Recently, the ENCODE Project has released an encyclopedia of CREs (SCREEN), which leverages available chromatin modifications, chromatin immunoprecipitation sequencing (ChIP-Seq) binding experiments, and expression quantitative trait loci (eQTLs) to annotate putative regulatory elements.(11, 12) Other attempts at annotating the regulatory genome were performed with genome-wide Cas9 editing screen (13), and more narrowly with targeted Massively Parallel Reporter Assays.(2) Measuring CRE activity using self-transcribing active regulatory region sequencing (STARR-Seq) has proved to be adaptable and sensitive, making it a powerful assay in the functional genomic toolkit.(14) However, despite this work, CRE imputation is limited in functional predictive power, and screening approaches, including STARR-Seq, largely overlook negative regulatory elements (NREs), such as silencers, attenuators, and insulators.(15)

Enhancer functional screening has been the subject of numerous genome-wide investigations, including in some immune cell populations.(16–19) NREs, however, are historically the subjects of individual experimentation. Notably, NREs are uniquely important during the development of some immune cell populations, in which they suppress CD4 expression during thymic development.(20) Only recently have silencers been identified through novel genome-wide screening methods.(2, 3, 16) Such screens are limited by the use of a synthetic library, or the ability to identify only NREs and not enhancers. One recent report suggests that ATAC-STARR is able to detect both activating and repressing fragments, but this study was performed in a cell line and does not include functional validation.(1) There remains a need to functionally assess the regulatory potential of CREs in cell types of biomedical interest, particularly in primary, non-transformed, human cells.

CD4+ T cells are adaptive immune cells with important roles in human health and disease. It is known that CD4+ T cells are transcriptionally plastic, responding to cytokine stimuli with diverse and distinct transcriptional programs.(21) These large transcriptional changes are accompanied by precise opening and closing of chromatin; however, the functional status of these putative CREs remains unknown.(10) A limited number of studies have attempted to define CREs in CD4+ T cells, including using a synthetically constructed massively parallel reporter assay (MPRA) to identify small pathogenic single-nucleotide polymorphisms (SNPs)(22) and using pseudo genome-wide assays like CapSTARR-Seq (DNA hypersensitive site enrichment with capture) in P5424 murine thymocytes.(23) Others have proposed a CD4+ subtype specific enhancer atlas based on histone 3, lysine 27 acetylation (H3K27ac) and histone 3, lysine 4 monomethylation (H3K4me1) distribution.(24) Thus, there is an active need to comprehensively profile the function of the non-coding genome of relevant human cells, in particular human CD4+ T cells.

We implemented a STARR-Seq–based screen in human resting total CD4+ T cells using a non-integrating lentiviral transduction system. Our screen identifies nearly 5000 functional enhancers and 5000 functional NREs from a library of open chromatin. This assay is the first example of a genome-wide assay examining the accessible chromatin elements in human primary cells. Herein, we demonstrate that a modified STARR-Seq screening method can identify both enhancer and NREs with high specificity as validated by luciferase assays. We find that the STARR-Seq enhancers and NREs are marked with distinct profiles of histone modifications, including both canonically activating and repressive histone marks. Interestingly, enhancers and NREs display distinct nucleosome positioning in their endogenous locations, and provide regulation of target genes via chromatin looping to target promoters. We provide supporting evidence that NREs may function as enhancers in other cell types, whereas functional enhancers are largely specific to hematopoietic lineages. We also provide a catalogue of transcription factors that may regulate enhancers and NREs in CD4+ T cells, providing hitherto known and unknown factors for future study.

## Materials and Methods

### Reagents

Reagents:
| Catalogue Number | Kit | Manufacturer | Location |
| --- | --- | --- | --- |
| 11346D | Human Untouched CD4+ Isolation Kit | Invitrogen | Waltham, MA, USA |
| A1048501 | CTS Optimizer | Gibco | Waltham, MA, USA |
| G7513 | L-Glutamine Supplement | Sigma-Aldrich | Burlington, MA, USA |
| P4333 | Penicillin-Streptomycin Solution | Sigma-Aldrich | Burlington, MA, USA |
| C3040H | Stable E. Coli | New England Biolabs | Ipswich, MA, USA |
| E0554S | Q5 Site-Directed Mutagenesis | New England Biolabs | Ipswich, MA, USA |
| D4203 | Plasmid Endo-Free Maxi-Prep | Zymo | Irvine, CA, USA |
| 28206 | MinElute Reaction Cleanup Kit | Qiagen | Germantown, MD, USA |
| 28104 | PCR Cleanup Kit | Qiagen | Germantown, MD, USA |
| M0544S | NEBNext Ultra-II Q5 PCR Master Mix | New England Biolabs | Ipswich, MA, USA |
| T1020L | Gel Purification Kit | New England Biolabs | Ipswich, MA, USA |
| E5520S | NEBuilder HiFi DNA Assembly Cloning | New England Biolabs | Ipswich, MA, USA |
| 75144 | RNeasy Midi | Qiagen | Germantown, MD, USA |
| T2040L | Monarch RNA Cleanup Kit | New England Biolabs | Ipswich, MA, USA |
| 18090010 | SuperScript IV | Invitrogen | Waltham, MA, USA |
| 28206 | Reaction Cleanup Kit | Qiagen | Germantown, MD, USA |
| C640003 | MegaX DH10B T1-R Electrocompetent Cells | Invitrogen | Waltham, MA, USA |
| 165-2089 | Gene Pulser Cuvette 0.1cm | BioRad | Hercules, CA, USA |
| BP1426 | LB Broth, Miller | Fisher | Fair Lawn, NJ, USA |
| 3471 | Ultra Low Bind 6-Well Plate | Corning | Glendale, AZ, USA |
| 11995-065 | Waltham, MA USA | Gibco | Waltham, MA, USA |
| S11150 | Fetal Bovine Serum | R&D Systems | Flowery Branch, GA, USA |
| 23966 | Polyethylenimine MW~25,000 (PEI) | Polysciences, Inc | Warrington, PA USA |
| 205548 | Caffeine | Millipore | Burlington, MA, USA |
| TR1003G | Polybrene | MilliporeSigma | Burlington, MA, USA |
| N1610 | Nano Glo Dual Luciferase Assay | Promega | Madison, WI, USA |
| 3912 | Solid White 96-Well Plate | Corning | Glendale, AZ, USA |
| MPK10025 | Neon Electroporation Kit, 100 µL Tips | Invitrogen | Waltham, MA, USA |

### Biologic Resources

Biologic Resources:
| Catalogue Number | Reagent | Manufacturer | Location |
| --- | --- | --- | --- |
| 63586 | psPAX2-D64V | Addgene | Watertown, MA, USA |
| 133996 | pLAI-Env | Addgene | Watertown, MA, USA |
| 45968 | pCMV-Enh-nLUC | Addgene | Watertown, MA, USA |
| 6251 | pHRL-TK | Promega | Madison, WI, USA |
| 8441 | pGL4.26 | Promega | Madison, WI, USA |
| NA | HEK-293T | ATCC | Manassas, VA, USA |
| NA | Leukoreduction Filters | Hoxworth Blood Center | Cincinnati, OH, USA |

### Human CD4+ Isolation

Peripheral blood mononuclear cells (PBMC) were isolated from Lymphocyte Reduction Filters obtained from deidentified male donors at Hoxworth Blood Center at the University of Cincinnati. CD4+ T cells were isolated by magnetic negative selection using Untouched Human CD4+ Isolation Kit (Invitrogen 11346D) per manufacturer’s recommendations. CD4+ T cells were cultured in Optimizer CTS (Gibco #A1048501) with 2 mM L-Glutamine supplement (Sigma # G7513) and 1X Penicillin-Streptomycin (Sigma # P4333) in incubators maintained at 37°C with 5% CO2.

### pORI-STARR-LV Construction

The lentivirus backbone plasmid was a gift from Kazuhiro Oka (Addgene plasmid #72263, RRID:Addgene_72263). This vector was digested with BspDI and KpnI and gel purified. A custom double-stranded (ds)DNA gBlock containing the origin of replication (ORI) and PolyA sequence was obtained from IDT (Supplemental Table 1), modeling the hORI-STARR screening vector generated by the Stark laboratory (Addgene plasmid #99296)(25). Immediately following cloning, the original ORI was excised using a Site Directed Mutagenesis kit (NEB E0554S). The vector was transformed (NEB C3040H) and isolated using Endotoxin Free Maxi-Prep (Zymo D4203).

### Lenti-STARR Library Preparation

The STARR-Seq inset library was created according to the ATAC-STARR-Seq approach.(17) Briefly, OMNI ATAC-Seq was performed as previously described with 320,000 human CD4+ T cells (in total) in batches of 100,000 cells per reaction (NEB #M0544S).(26) The ATAC-Seq reaction products were cleaned up using the Qiagen Reaction Cleanup Kit (Qiagen #28206) and eluted in 12 μL per 100,000 cells transposed. Each resultant elution was amplified in 50 μL with custom primers (Supplemental Table 1) for 10 total cycles, as described previously.(17) The amplified library was purified using a PCR Cleanup Kit (Qiagen #28104) and size selected for 150-500–bp fragments on a 2% agarose gel, using 300 mg per column (NEB #T1020L). The eluted ATAC-Seq library was cloned into AgeI- and SalI-digested pLenti-STARR using a 3:1 molar ratio (insert:backbone) in a total reaction volume of 100 μL using the NEBuilder HIFI DNA Assembly Kit (NEB #E5520S). The reaction was concentrated to a total of 20 μL in a Reaction Cleanup Kit (Qiagen #28206). The cloned ATAC-Lenti-STARR was electroporated into MegaX DH10B cells (Invitrogen #C6400003) using 10 μL library per 100 μL cells [2.0 kV, 200 Ω, 25 μF] in 20 μL electroporation reactions in 0.1-cm cuvettes (BioRad #165-2089) on a Harvard Apparatus (ECM Model 630). Immediately following electroporation, 950 μL of prewarmed recovery media was added, and cells were cultured for one hour at 37°C. All electroporation reactions were pooled to 1 L sterile Luria Broth (Fisher #BP1426) and grown in 0.2 L per 2 L flask for 8 h at 37°C, 300 RPM. Each 200 mL of culture was purified in Endotoxin Free MaxiPrep Reactions (Zymo #D4203) and eluted in 0.4 mL elution buffer per column.

### Lentivirus Preparation

HEK-293T cells were cultured in 15-cm^2^ dishes in Dulbecco’s Modified Eagle’s Medium (DMEM) (Gibco #11995-065) with 10% fetal bovine serum (FBS) (R&D Systems #S11150). Cells were grown to 70% confluency, then DMEM was replaced two hours before transfection. Each 15-cm^2^ plate was transfected with a total of 21 μg of plasmid DNA (1:2:1 ratio psPAX2-D64V:pLenti-STARR:pLAI-Env) [psPAX-D64V (Addgene #63586), and pLAI-HIV (Addgene #133996)]. DNA was mixed with 4.25X polyethylenimine (PEI) (Polysciences #23966) in at total of 2 mL Dulbecco’s Phosphate-Buffered Saline (DPBS per plate), left to rest for complex formation for 10 minutes at 22°C, and then added dropwise to the cell culture while swirling. Ten hours after transfection, the cells were washed once with DPBS and then cultured in DMEM with 10% FBS and 2 mM caffeine (Millipore #205548). Forty-eight hours after transfection, the media was collected and held at 4°C. Seventy-two hours after transfection, the media was collected and pooled. The viral supernatant was sterilely filtered through a 0.45-μM PES filter, loaded into ultracentrifuge tubes (Beckman #344058), and spun for 2.5h at 4°C and 35,000 g. Each viral pellet was dissolved in 200 μL DPBS and left overnight at 4°C to resuspend. Concentrated lentivirus was resuspended, filtered through a 0.45-μM PES filter, and snap frozen in an ethanol/dry-ice bath at −80°C until use.

### Lenti-STARR Transduction

Freshly isolated human (h)CD4+ T cells were suspended to 6.6E6 cells per mL in CTS Optimizer and then combined with 0.33 mL concentrated lentivirus per mL of cells. Cells were spinfected in low-bind 6-well plates (Corning #3471) with 7 μg/mL polybrene (MilliporeSigma #TR1003G) at 950 *g* and 32°C for 2 hours. Immediately following spinfection, cells were gently resuspended and cultured for 2 hours at 37°C, 5% CO_2_. Cells were then pooled, centrifuged, and resuspended in Optimizer CTS for 24 hours of culture. Cells were treated with 100 units of DNaseI (NEB M0303S) for 30 minutes at 37°C for dead cell removal.

### Lenti-STARR Library Construction

Total RNA was purified from transduced cells (Qiagen #75144) and concentrated 7.5X using an RNA Cleanup Kit (NEB #T2040L). RNA was reverse transcribed according to manufacturer guidelines with STARR-Seq transcript–specific primers (Supplemental Table 1) with SuperScript IV (Invitrogen #18090010), using twice the recommended RNA input. cDNA was purified using a Reaction Cleanup Kit (Qiagen #28206) and amplified using custom STARR-Seq transcript–specific primers (Supplemental Table 1) as previously described.(17) Diluted Lenti-STARR-Seq plasmid ‘input’ control was amplified using an input concentration that yielded the same amplification cycle number as the STARR library for hCD4+ Donor I. PCR amplified library was purified using PCR cleanup columns (Qiagen #28206) and quantified for PE-150bp sequencing (Novogene).

### Alignment and Next-Generation Sequencing Processing

Samples were aligned to hg19 using SciDAP.com with the ChIP-PE processing pipeline (https://github.com/datirium/workflows/blob/master/workflows/trim-chipseq-pe.cwl).(27) Briefly, reads were trimmed and aligned to hg19, and bam files were produced. All mitochondrial reads were removed. The per duplication count of each unique read for each biologic donor was fitted to the negative binomial distribution for each technical replicate (input plasmid library). Duplicate reads exceeding the 75th percentile were removed from each sample by MACS2 to reduce false discovery of enhancers resulting from PCR duplication. Enhancers were called using MACS2 in which the plasmid reads were used as a control. Enhancers were retained if they i) had a MACS2 logP value greater 50 *and* ii) intersected with a MACS2 peak called from input plasmid reads alone. NREs were annotated with Fast-NR again using input plasmid reads as a control, with the following parameters: cosine similarity 0.5, adjusted p-value less than 0.05.(28) Significant NREs were retained if they i) had a FAST-NR–adjusted logP value greater than 30 *and* ii) intersected with a MACS2 peak called from input plasmid reads alone.

### Heatmaps

All heatmaps were constructed using deeptools.(29)

### Nucleosome Position Imputation

Reads mapped to the input plasmid to ATAC-STARR were sequenced as described above. Bam files were processed with NucleoATAC to produce nucleosome occupancy scores and converted to a .bigWig for heatmap visualization.(30)

### Luciferase Assays

Putative regulatory elements were randomly selected from STARR-Seq functional enhancers and NREs. The sequences for functional testing were selected on the basis of fragment distribution in the control ATAC library and sought to include the input ATAC-Seq peak center. gBlocks or eBlocks were synthesized (IDT) ranging from 150-400 bp in length and cloned into pGL4.26 (Promega #8441) for enhancer screening or CMV-Enh-Luc (Addgene #45968) for NREs screening (All tested sequences included in Supplemental Table 1). Total human CD4+ T cells were purified as described above. Each luciferase experiment was performed using two donors and in technical transfection duplicate. Each transfection reaction consists of 20 μg of testing vector and 500 ng of Renilla-expressing phRL-TK (Promega #6251) mixed with 3.5E6 cells in buffer T using 100 μL Neon Tips (Invitrogen #MPK10025) [2200 V, 20-ms pulse width, 1X pulse]. Cells were cultured for 24 hours, collected, and suspended in 60 μL DPBS for Nano-Glo Dual Luciferase Assay, according to the manufacturer’s instructions (Promega # N1610). Absorbances were read using the Dual-Nano-Glo protocol with a GloMax plate reader (Promega #GM3000) in flat-bottom, opaque, white, 96-well plates (Corning #3912). Absorbances were averaged across the technical transfection duplicates. Control transfection (empty vector) was performed for each CD4+ T cell donor and plasmid condition. Error is presented as standard deviation, and statistical significance is indicated if control and experimental [1.96*SD] ranges were non-overlapping (95% confidence interval).

### RNA Expression Analysis

RNA expression values [transcripts per million (TPM)] were downloaded from PolyA RNA-Seq, performed in total CD4+ T cells (ENCSR545MEZ). The Nearest TSS of STARR-Seq CRE were annotated by Homer and retained if within 10 kb of the CRE.(31) Genes were assigned to the following bins: targets of enhancers alone, NREs alone, both enhancers and NREs, or ATAC site association alone. CRE and target promoter assignments were annotated using Promoter-Capture Hi-C performed in total CD4+ T cells.(32) CRE were intersected (bedtools intersect), and only high-confidence chromatin interactions were retained (CHiCAGO score greater than 5).(33) Statistical differences in target gene expression between CRE groups were assessed with a non-parametric, Kruskal-Wallis two-sided test with Holm adjustment for multiple comparisons. Significant differences in TPM were annotated if the adjusted p-value were less than 0.01. A multivariate linear model explaining TPM by the number of CRE:Promoter contacts was used to test for significance between gene expression and the number of enhancers, NREs, or only ATAC-Seq peak contacts. The Beta slope coefficients of each independent variable in the linear model (TPM ~ NREs Contact Number + Enhancer Contact Number + ATAC-Seq Peak contact number + b) are presented along with standard error and annotated as significant if the p-value were less than 0.001.

### Statistical Analysis

All statistical tests were performed in R and detailed in the appropriate methods sections and figure legends.

### Data Availability

All generated datasets from Lenti-STARR have been deposited in GEO under GSE217535. Other datasets utilized in this manuscript can be found listed in the supplemental table ‘Utilized Datasets’.

## Results

### Lenti-STARR-Seq in Primary Human CD4+ T Cells

We adapted a screening vector from the promoter-less STARR-Seq vector, which uses a bacterial origin of replication (ORI) as a cryptic promoter to initiate transcription (**1A**).(25) This STARR-Seq screening vector was further cloned into a lentiviral backbone for viral packaging. An open chromatin library was prepared from resting total human CD4+ T cells from a single donor in 16X reactions with 100,000 cells each to ensure sufficient library diversity. The OMNI-ATAC protocol was employed to reduce the number of mitochondrial reads without the need for negative selection.(26) ATAC-Seq library fragments were gel purified (150-500 bp including adapters), cloned into pLenti-STARR **(1B)** using NEBuilder HiFi cloning assembly, and amplified in bacteria. We elected to use lentiviral particles with an HIV-1 envelope to allow for transduction of resting CD4+ cells.(34, 35) A lentivirus packaging vector with mutated integrase, psPAX-D64V, was used during lentiviral packaging to ensure that the STARR-Seq plasmid stays episomal, thus avoiding positional regulatory effects.(35) The Lenti-STARR-Seq library was transduced into human CD4+ T cells isolated from four separate adult, healthy donors. At least 50 million CD4+ cells were transduced per donor to ensure that the full library diversity is assessed in each biological replicate. Twenty-four hours after transduction, total RNA was harvested, reverse transcribed with a STARR transcript specific primer, PCR amplified, and next-generation sequenced. An input control library was amplified from the pLenti-STARR-Seq library.

Enhancers were called using MACS2, with the input plasmid library used as a control. Only enhancers overlapping peaks called from the input control were considered for downstream analysis. Although the weaker ATAC sites may function as enhancers within a STARR-Seq assay, we reason that the lower accessibility makes them less likely to be functional in their native genomic context. NREs, likely including silencers and insulator sequences, were detected using the FAST-NR algorithm, which leverages both STARR-Seq RNA repression and coverage dissimilarity to detect significant repression.(28) Significant NREs were also filtered to only include sites that overlap MACS peaks called from the input plasmid library alone.

STARR-Seq functional enhancers demonstrate strong central STARR RNA production, whereas NREs are depleted in the STARR-Seq library relative to all peaks from input plasmid control (**1C**). Although the Lenti-STARR screening library is shared between donors, STARR RNA regulation appears consistent across the hCD4+ donors (**1D**) as demonstrated by high Pearson correlation coefficients of read counts across peaks.

### Enhancer and NRE Element Validation

After identifying functional CREs by STARR-Seq, we sought to validate their function by traditional luciferase assays (**2A)**. Functional NREs in STARR-Seq were cloned upstream of a strong promoter (pCMV-ENH-LUC) and transfected into resting hCD4+ T cells. Luciferase activity was calculated relative to a Renilla luciferase (pRL-TK) transfection control. Statistical significance is calculated by comparing 95% confidence intervals [mean +/−1.96 * SD]; if this confidence interval was not overlapping signal from control (donor specific), then repression was considered significant. We find that the majority of putative NREs were functional in traditional luciferase assays, across varying FAST-NR significance and activity (fold change) levels, and across a wide array of endogenous genomic locations (intron, exon, near promoter, intergenic). Some fragments displayed in Figure 2 were tested in luciferase assays despite not meeting strict filtering requirements.

**Figure 1:**
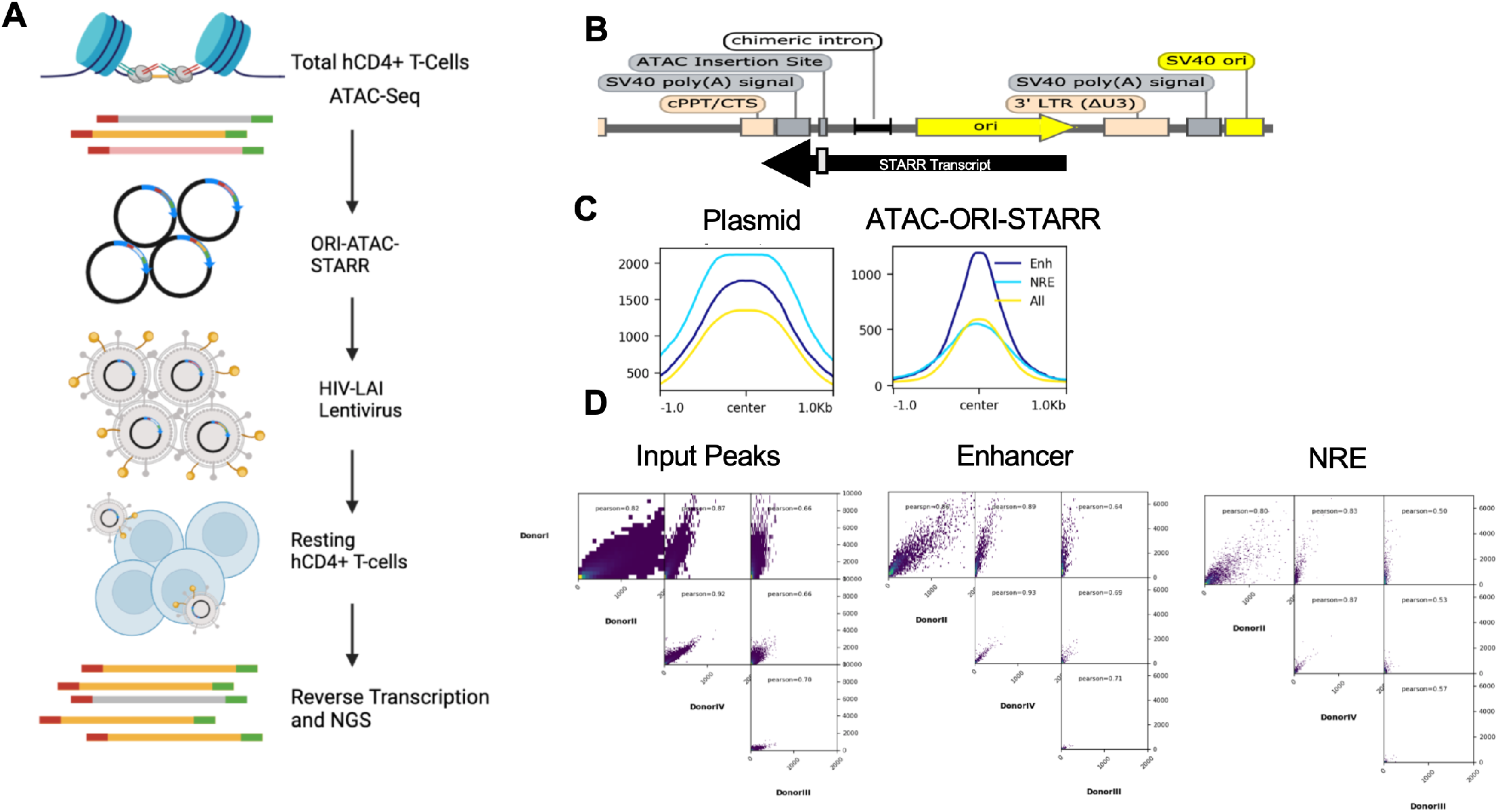
Lenti-STARR Approach. **A)** Putative regulatory sequences enriched by ATAC-Seq are used as input into Lenti-STARR-Seq. ATAC-Seq was performed on total human CD4+ T cells from peripheral blood obtained from healthy adult donors. The ATAC-Seq library is cloned into a promoter-less STARR-Seq screening vector that uses a bacterial origin of replication as a cryptic promoter.^1^ The screening library is packaged in an HIV-enveloped non-integrating lentivirus, and transduced into resting CD4+ T cells. After 24 hours in culture, STARR-Seq RNA is collected, reverse-transcribed using a plasmid-specific primer, PCR-amplified, and prepared for next-generation sequencing (NGS). **B)** The Lenti-STARR-Seq screening vector carries an ATAC insertion site downstream of chimeric intron. **C)** Enhancers are called with MACS2, comparing input plasmid reads to STARR-Seq–obtained RNA reads.(74) Negative regulatory elements (NREs) are called using FAST-NR.(28) Mean tag density is displayed, compared to ‘All’ open peaks from input. **D)** Pearson correlation of tags across either Input Peaks, Enhancers, or NREs for each STARR-Seq human CD4+ donor (n=4).

**Figure 2:**
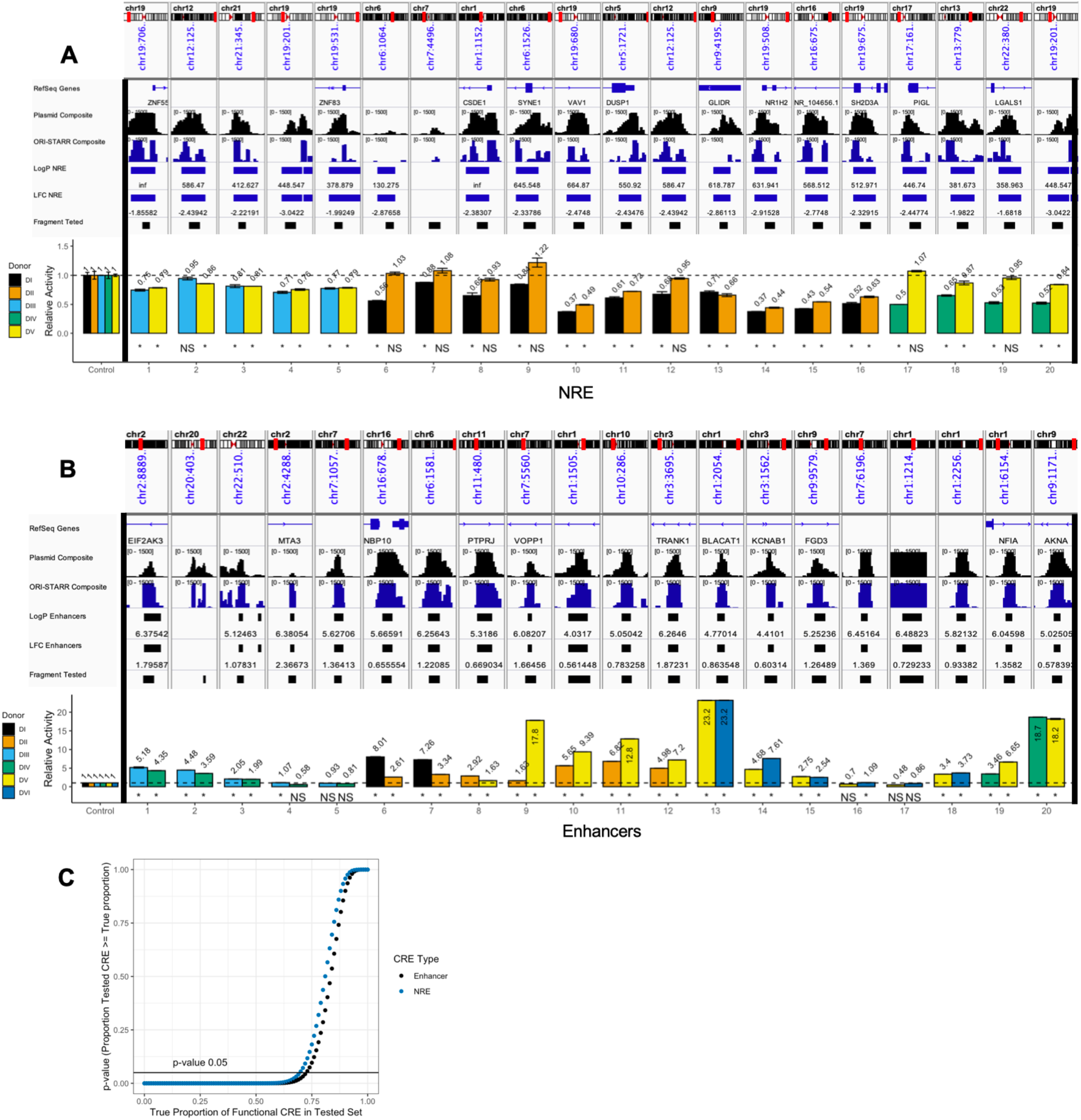
CRE Validation by Luciferase Assays. MACS2 Log2FC (for enhancers) and FAST-NR Log2FC (for NREs) are displayed in the genome browser above. Fragment Tested represents the DNA sequence cloned for luciferase testing. (*) Non-overlapping 95% Confidence Intervals. Error represented as Mean ± SD. **A)** Twenty putative NREs are tested using CMV-Enhancer-LUC in DualGLO Luciferase assays (Promega). Two independent human CD4+ Donors were used for each CRE tested, with two technical replicates each. Relative Activity compares NRE activity to empty-vector CMV-Enhancer-LUC control activity. **B)** Twenty putative enhancers are tested using pGL4.26 (minP promoter) in DualGLO Luciferase assays as described above. **C)** Binomial exact test trials to estimate the true fraction of tested CREs that are functional in luciferase assays. P-value indicates the likelihood that the observed proportion of luciferase active fragments is greater than the theoretical proportion on the x-axis.

STARR-Seq–identified enhancers were cloned upstream of the minimal promoter (pGL4.26) luciferase-expressing vector. Enhancers demonstrated strong luciferase activity, which was sometimes donor dependent. (**2B**). Similar to NREs, functional enhancers identified by STARR-Seq came from across various genomic locations (intron, intergenic, exon, near promoter).

We next estimated the true proportion of enhancers and NREs that are functional within the original screening library. The true proportion of CRE activity was estimated from the success rate of luciferase validation using binomial exact tests (**2C**). It has been suggested that enhancers comprise between ~10% (36) and 35% (37) of accessible chromatin sites. We found that 17% of ATAC-Seq peaks in CD4+ T cells exhibited statistically significant enhancer activity by STARR. The estimated true proportion of these enhancers that are also luciferase active was significantly greater than 75% (p-value < 0.05) (**2C**), with a maximum-likelihood estimated at nearly 80%. STARR functional NREs comprised nearly 17% of the accessible genome in CD4+ T cells and also yielded an estimated true proportion of functional NREs of nearly 75% by luciferase validation. Although it is unknown what fraction of the accessible chromatin landscape comprises NREs, we observed overall that STARR-Seq performed similarly for enhancers and NREs, but may marginally outperform in detecting functional enhancers over functional NREs.

### STARR-Seq CREs Regulate Transcription Within Their Endogenous Location

As expected, STARR-Seq enhancers and NREs display a distinct relationship with RNA transcriptional initiation and RNA polymerase. Assays, including Precision Run-On Nuclear Sequencing (PRO-Seq), capture nascent RNA production (bidirectional non-polyadenylated RNA) characteristic of active enhancer elements. (38, 39) (40) Shown in Figure (**3A**), we find functional enhancers in STARR-Seq displayed strong central enrichment of nascent RNA production in PRO-Seq, whereas NREs were centrally depleted of such transcripts. Intriguingly, there was enrichment of the PRO-Seq signal within 100-200 bp around NREs, suggesting that enhancers may be located in surrounding regions. This suggests that although NREs are located in the genomic areas generally permissive to eRNA transcription, transcriptional initiation is centrally repressed. Although the NREs may include silencers and insulators, the clear repression observed in the PRO-Seq signal suggests that this group of CRE is distinctly less transcriptionally permissive. NREs and enhancers both demonstrated RNA Polymerase II binding, though NREs exhibited weaker enrichment of S5-Phosphorylated PolII. These findings suggest that PolII may be recruited to enhancer elements bordering NREs but is less likely to be functionally active (**3B**).

We next sought to characterize the effect of enhancers and NREs on expression of their target genes. First, we attempted to assign target genes to CREs on the basis of distance. Each CRE was assigned to the nearest TSS if within 10 kb (**S7**). We found that genes assigned enhancers or both enhancers and NREs had significantly higher TPM than genes associated with only open peaks from input. However, the genes nearest NREs also had higher expression than did the genes associated with only open peaks from input. We posit that the close physical assortment of enhancers and NREs may result in false nearest TSS assignment because of intervening NREs (e.g., insulators) and complex 3D genomic architecture. Another potential explanation is that some of the regulatory elements shown to be functional in STARR-Seq or luciferase assays were not active in their endogenous epigenomic context (see Discussion).

Although assignment of genes to CREs by distance was used traditionally, multiple reports suggest that both enhancers and NREs loop from long distances to activate or repress their target promoters.(3, 41) We assigned CRE gene targets leveraging Promoter Capture HiC (PC-HiC) performed in total CD4+ T cells.(32) Genes that are physically contacted by NREs are significantly repressed compared to those contacted by enhancers (**3C**). Genes that are the targets of both enhancers and NREs or enhancers alone had significantly higher expression than genes that were targets of non-annotated ATAC-Seq peaks. We assessed whether the regulation by CRE was dose-dependent on the number of CREs that target each promoter. RNA expression was predicted by the number of CRE contacts using linear regression. We found that the number of NREs that contacted a gene was a significant negative predictor of gene expression (ß_NREs_: −19.97 +/−6.48) (**3D, 3E**). The number of enhancers that contacted a gene was a positive predictor of expression, but did not reach statistical significance (ß_ENH_: 2.51 +/−2.06). It is well known that enhancers are often cryptic and redundant (42, 43), which supports our finding that genes with at least one enhancer association displayed significantly higher expression in Figure 3C, but that the presence of additional enhancers was not associated with a further increase in expression. We cannot rule out that unannotated CRE are functional regulators of expression, as non-STARR functional ATAC-Seq peaks demonstrate significant positive influence on expression of target genes (**3E)**.

**Figure 3:**
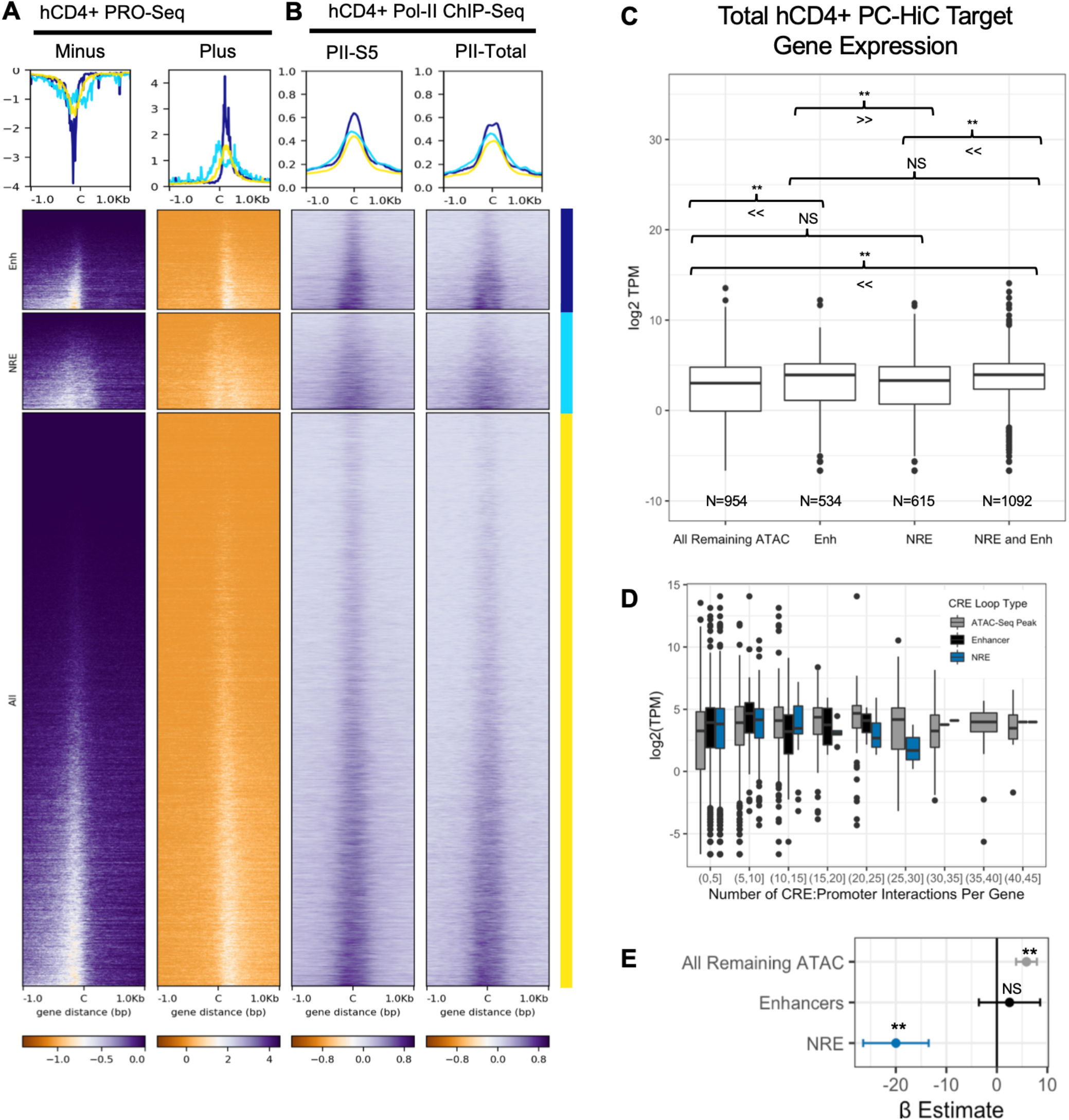
CRE Regulation of Transcription. **A)** PRO-Seq tags in hCD4+ T cells (GSE66031) at STARR-Seq Enhancers, NREs, and ‘All’ open peaks from input plasmid library. **B)** ChIP-Seq experiments of RNA Polymerase II Phospho-S5 and total RNA Polymerase II conducted in human CD4+ T cells from Barski *et al*. 2007 **C)** Chromatin looping targets of CRE to promoter assigned from Javierre 2016, with TPM from hCD4+ PolyA RNA-Seq (ENCSR545MEZ). (** P-value < 0.01 by non-parametric Kruskal-Wallis two-sided test with Holm adjustment for multiple comparisons). **D)** Number of CRE:Promoter PC-HiC contacts to each target gene, with TPM from hCD4+ PolyA RNA-Seq (ENCSR545MEZ). **E)** Multivariate linear model explaining TPM (ENCSR545MEZ) by number of CRE:Promoter interactions to each promoter [TPM ~ ß_ATAC_ + ß_ENH_ + ß_NREs_]. ß slope coefficients ± Standard Error displayed. (** P-value < 0.001)

**Figure 4:**
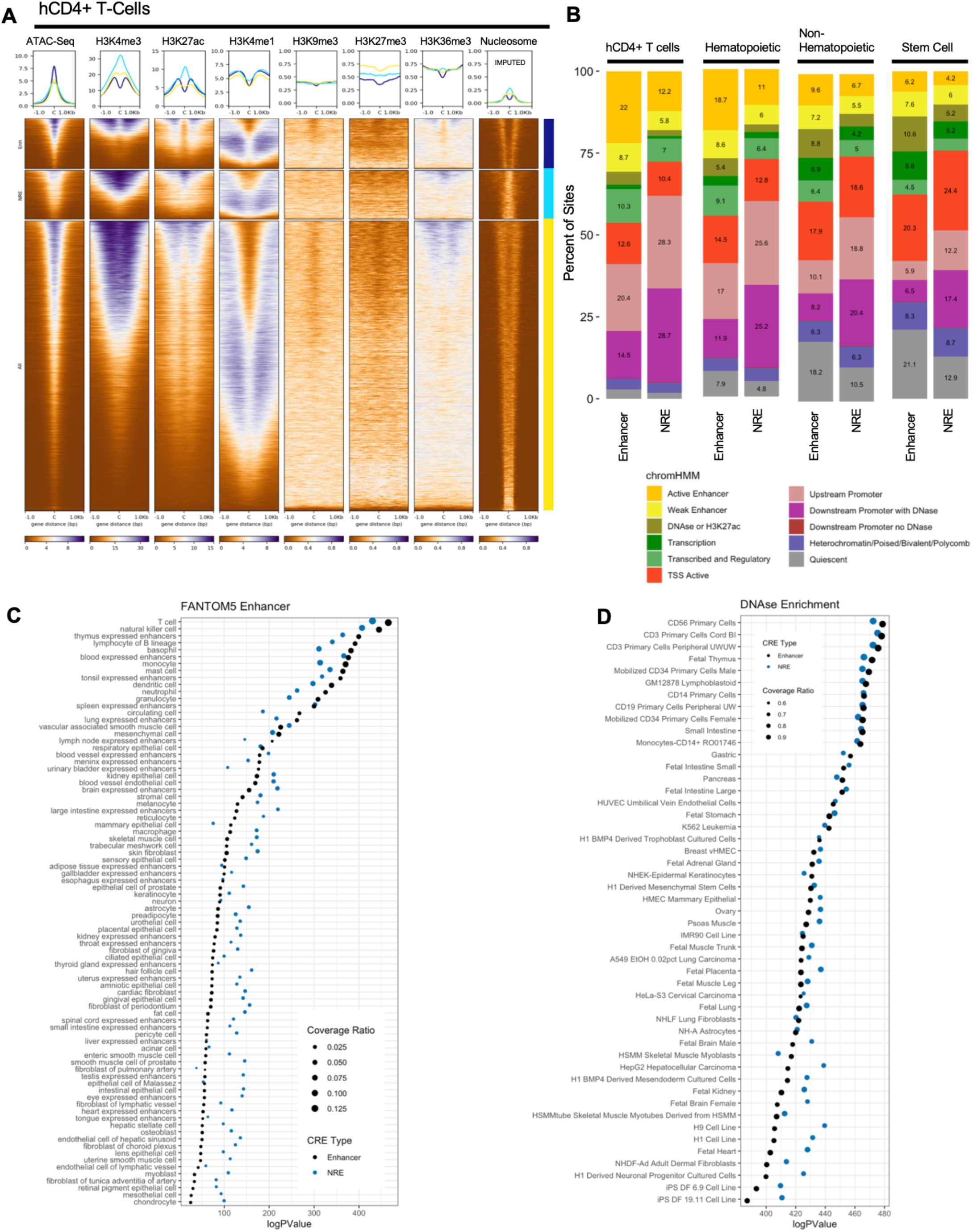
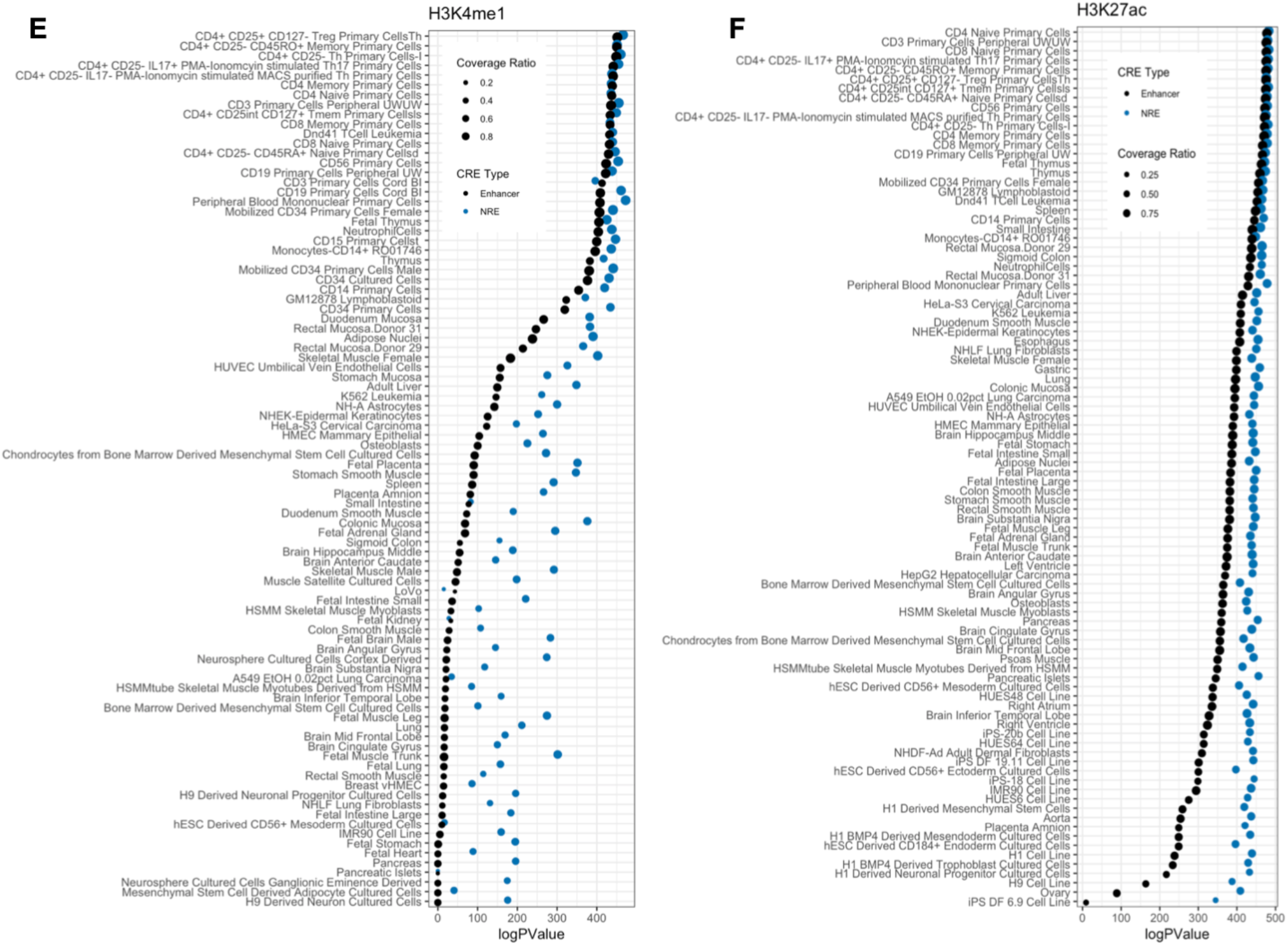
Chromatin Environment of Functional Regulatory Elements. **A)** Tag density plots of mean intensity and heatmaps of ChIP-Seq or ATAC-Seq performed in human CD4+ T cells plotted against STARR-Seq– identified Enhancers, NREs, and ‘All’ open peaks from input. (ENCODE). Nucleosome location in resting CD4+ T cells imputed from NucleoATAC. Tag density of mean signal intensity displayed for Histone enrichment, and of imputed nucleosome position. **B)** Proportion of functional CD4+ STARR-Seq Enhancers and NREs within ChromHMM Segmentation of 25 imputed groups. Other cell groups displayed as percent of total. **C**) RELI enrichment logP-value of available enhancer predictions in FANTOM5 database. Coverage ratio is the proportion of target sites overlapping all sites. **D**) RELI enrichment logP-value of STARR-Seq identified elements across available DNase-Seq experiments, **E**) H4K1me1 experiments, and **F)** H3K27ac experiments.

### Lenti-STARR NREs and Enhancers Exhibit Distinct Patterns of Chromatin Modifications

We next sought to compare histone modification landscape across functional enhancers and NREs identified by STARR-Seq. Previous functional silencer screens report enrichment with chromatin modifications H3K27me3, H3K9ac, and H3K79me2, which were also enriched at the functional NREs identified in this study (**S1**).(1, 3, 44, 45) Enhancers classically display enrichment with H3K27ac, H3K4me1, and sometimes H3K4me3, which we also observe at our STARR functional enhancers as well. Reports of other functional silencers from screens of open chromatin find some enrichment for marks of heterochromatin(3, 45). Functional NREs identified in our assay were enriched centrally for H3K27me3 and H3K9me3, classical heterochromatin marks, whereas functional enhancers displayed characteristic central depletion of these marks (**4A**). A mark related to transcriptional elongation across gene bodies, H3K36me3, was also central enriched at STARR-Seq NREs and depleted at enhancers, as was seen previously(3). We found that the traditionally active histone marks H3K27ac and H3K4me3 demonstrated divergent enrichment patterns at functional enhancers and NREs. H3K4me1, another mark of active enhancers, appeared similarly enriched at both NREs and enhancers. The ATAC-Seq tag-density plot highlighted endogenous differences in chromatin structure, in which NREs tended to be *less centrally accessible*. The distribution of H3K27ac and H3K4me3 histone ChIP-Seq tags suggested that NREs might be occupied by a nucleosome. To test this hypothesis, we imputed positions of nucleosomes from ATAC-Seq data using NucleoATAC (30). Functional NREs were occupied by nucleosomes (or possibly nucleosome-size protein complexes), whereas enhancers seemed to be located between nucleosomes.

To test where functional enhancers and NREs reside within previously imputed chromatin states, we overlapped these elements with genomic segments identified by ChromHMM imputation based on 25 histone marks (**4B**).(6) In CD4+ T cells, STARR-Seq functional enhancers were more likely to be segmented as enhancer subtypes, or as active TSS. NREs functional in STARR-Seq were more likely to be adjacent to promoters, particularly in downstream of TSS DNase accessible regions, which were likely intronic. NREs displayed weaker enrichment in putative enhancer classes than did STARR-Seq functional enhancers. Though STARR-Seq enhancers were most likely to be segmented as enhancers in imputation models in hCD4+ T cells themselves, we found that these enhancers were also predicted as enhancer groups across hematopoietic cells, suggesting that some CD4+ enhancers are functional within a broader class of hematopoietic cells. Neither STARR-Seq enhancers nor NREs frequently segmented in regions of heterochromatin or ZNF repeats in mature CD4+ or hematopoietic lineages; mature CD4+ T cells exhibited weak heterochromatin chromatin signatures amongst sites of open chromatin.(46) Intriguingly, functional NREs also appear within accessible and classically activated chromatin environments. We also observe that the percentage of both NREs and enhancer elements that were devoid of histone modifications (quiescent state) increased in non-hematopoietic cell types and stem cell lines, suggesting a progressive opening and activation of these CREs during development.

Both STARR-Seq identified enhancers and NREs were enriched within related cell types’ enhancer predictions in FANTOM5 (**4B**).(47) Consistent with other silencer reports, we found that NREs could convert into enhancers in unrelated cell types on the basis of both FANTOM5 enhancer annotations, and enhancer chromatin marks H3K4me1 and H3K27ac (**4D** and **4E**). Though we found that functional NREs are annotated as enhancers using their chromatin environment, additional experiments are needed to demonstrate that NREs are in fact cross-functional as enhancers in other cell types.

### Transcription Factor Binding Across Lenti-STARR Cis-Regulatory Elements

In order to identify putative transcription factors that interact with the regulatory elements that we identified, we examined the overlap between these elements and a collection of ChIP-Seq experiments from various cell types available in GEO database using the RELI algorithm.(48) Functional enhancers were likely to be bound by numerous, classic enhancer activating factors (**5A**). For example, CCAAT/enhancer binding protein zeta (CEBPZ) demonstrated stronger enrichment in enhancers versus NREs. We found that lymphoid enhancer binding factor 1 (LEF1) had strong enrichment at enhancers compared to NRE in epithelial carcinoma cells. LEF1 also has lymphocyte-specific functions: it promotes expression of some conventional and regulatory Th cell genes, while repressing CD8+ T cell–specific cytokine gene expression.(49, 50) Intriguingly, LEF1 enrichment was greater at NREs in embryonic stem cell ChIP-Seq experiments (**5B**). We observe that STARR-Seq functional enhancers are enriched with binding by some activation-inducible transcription factors: JUN (AP-1), FOS (AP-1), and RELA (NFκB) (**S2**). This suggests that at least some enhancers are capable of participating in, or become subsequently turned-on during CD4 activation. Constitutively expressed factors, including ETS family member FLI1, also were bound at enhancer elements. FLI is known to bind enhancers in T cells (51), and other immune cell subtypes [megakaryocytes(52)], and can induce T cell leukemia if overexpressed.(53) The FLI portion of FLI-Ewing’s Sarcoma fusion proteins was characterized as an activator of enhancers and activated reporters in luciferase assays.(54) Another putative enhancer binder that we identified was CD74, a cell surface marker which was recently shown to also have transcriptional activation activity.(55, 56)

We explored putative co-regulators of NREs by leveraging a catalogue of ChIP-Seq experiments, again performed in various cell types. Numerous histone modifiers are likely to play a role in NRE function; NREs demonstrated enrichment for lysine demethylases, including KDM2B and KDM4A, which are recruited to promote demethylation at H3K4, H3K9, and H3K36 residues. KDM4A demethylated both H3K36me3(57) and H3K9me3, and recruits NCOR (nuclear core repressor).(58) Other canonical repressors were also enriched in NREs, including members of polycomb repressor complex 1 (RNF2, RYBP) and polycomb repressor complex 2 (EZH2, SUZ12, EED). Another candidate for NRE function in CD4+ T cells is the known repressor PBXIP1 [(HPIP)], which inhibited PBX1 homeobox target activation(59) and is highly expressed in CD4+ T cells.

NREs may also be recruiters of chromatin remodelers. INO80 was recruited to NREs in human CD4+ T cells (**S2**), and is a chromatin remodeler recruited by YY1 and other TFs. We also found that CHD1 was recruited to NREs; another CHD family member CDH4 was also identified previously as binding to functional silencers in K562 cells.(3) Previous studies showed that CHD1 was capable of moving nucleosomes in response to binding of the canonical repressor LacI.(60) These current and prior findings support a model in which the accessibility changes and remodeling nucleosome position seen in **4A** may be a result of chromatin remodeler recruitment to NREs. Next, we conducted motif analysis to find transcription factor motifs enriched at CREs (**Figure 5, S3**). Similar to other reports, we found that STARR-Seq identified functional insulators in the negatively regulated fraction of screened sequences.(16) We observed that the CTCF motif was the second most enriched motif at NREs, behind only the ETS family motif, which was highly enriched across all open chromatin sites in CD4+ T cells. (**5C**). The function of CTCF is likely dual purpose; enhancers were also enriched for the CTCF paralogue, CTCFL (Boris), which is known to facilitate long-range enhancer and super-enhancer looping to target gene promoters.(61, 62) Other enriched motifs within both enhancers and NREs belonged to families with mixed activating and repressing function, including the RUNX and ETS families.(63, 64) Enhancers were also enriched for multiple activating transcriptional activators, including ATF and NFY, which frequently bound cell-specific enhancers(65), and SP2. We also found motif enrichment for transcription factors related to CD4 activation, suggesting that some of these enhancers are functional during CD4 activation (FOS).

**Figure 5:**
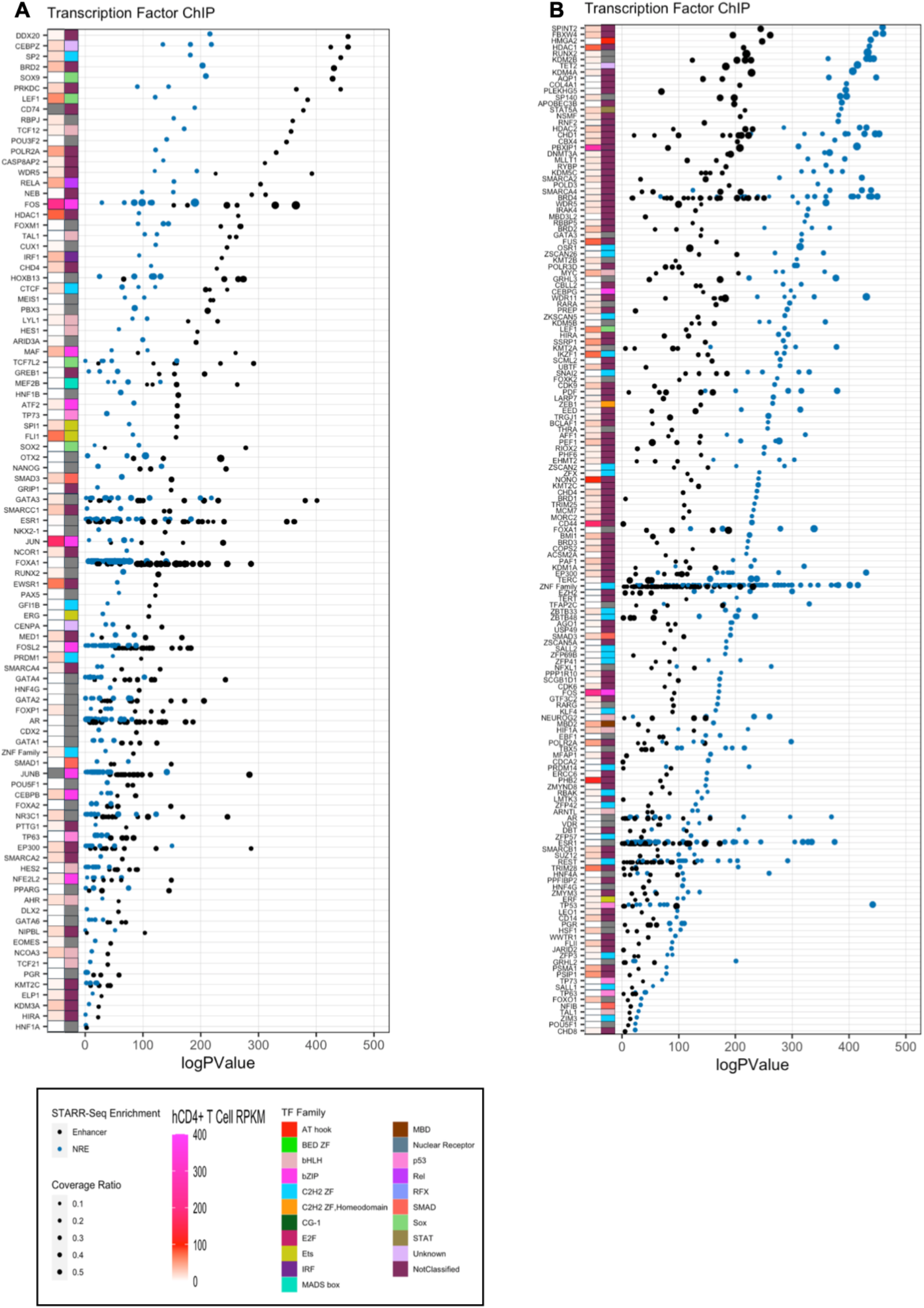
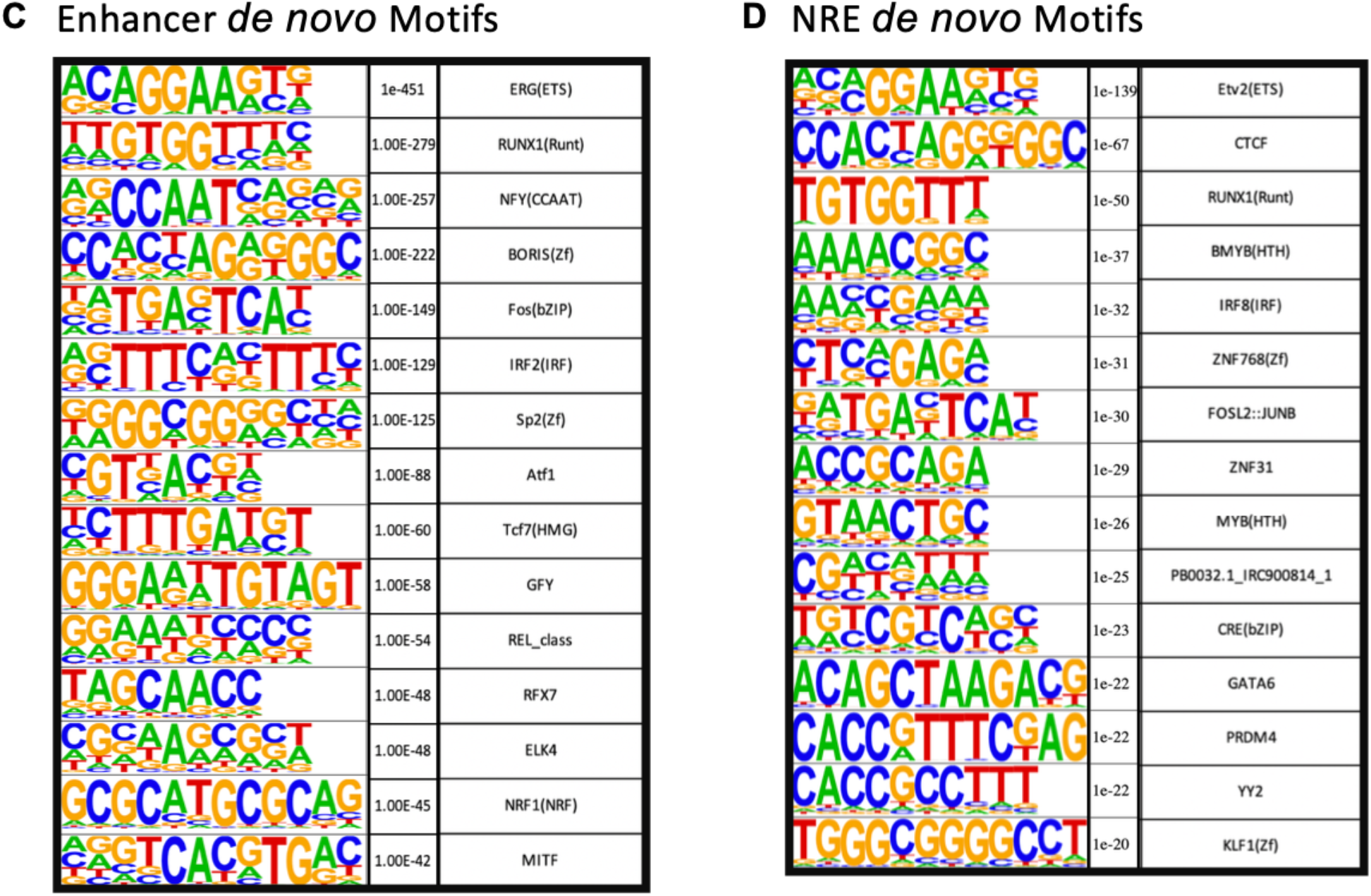
Transcription Factor Binding at CRE. RELI enrichment logP-value of CRE enrichment across available transcription factor ChIP-Seq experiments in GEO. Heatmaps of RNA expression [transcripts per million (TPM)] in resting human CD4+ T cells are displayed furthest left. Transcription factor family is displayed adjacent to RNA expression. **A)** The top 2500 ChIP-Seq experiments with strongest enrichment in STARR-Seq enhancers are displayed, whereas **B)** displays the top experiments with NRE enrichment. **C)** Motif enrichment performed in HOMER *de novo* motifs in Enhancers (predicted motif, p-value, and transcription factor name) and **D)** in NREs.

## Discussion

We describe a screen identifying functional enhancers and NREs from a library of open chromatin in resting human CD4+ T cells. Using a STARR-Seq approach, we identified and validated the function of CREs in human primary CD4+ T cells. We found that Lenti-STARR-Seq can identify enhancers and NREs, each with nearly 80% sensitivity, as verified by luciferase assays. We described the chromatin landscape of both enhancers and NREs, in which enhancers were marked with H3K27ac and H3K4me1. They were also enriched for binding of enhancer activating proteins (CEBPZ). Intriguingly, we found that NREs tend to have central nucleosome placement, unlike enhancers which are located between nucleosomes. NREs and enhancers also function to regulate gene expression via long-range chromatin interactions. In short, we report thousands of functional enhancers and NREs from a screen of CRE function in primary human CD4+ T cells.

Enhancers have long been known to bind activating factors, including P300 and CEBP(Z), among others. We also found in other cell types that CD4 enhancers could bind LEF, TCF, and novel, recently identified enhancer activators like CD74. Integration of chromatin accessibility and chromatin modification data across cell types suggests that many CD4 functional enhancers are broadly active across hematopoietic lineages. Indeed, many hematopoietic cells have overlapping repertoires of transcription factors that may bind these elements. Although many enhancers are likely shared between hematopoietic lineage cells, these findings invite further investigation to refine enhancer specificity within lymphoid cell subtypes, particularly during developmental transitions (e.g. CD8+CD4+ double-positive to CD4+CD8-single-positive cells), and activation events in immune cells. We also find that CD4+ enhancers are capable of being bound by inducible transcription factors following activation. (**S2**) Given that our experiments were performed in resting cells, it remains to be shown whether these enhancers are repurposed during activation by these inducible factors or whether lentiviral transduction may be weakly activating.

We conducted this STARR-Seq screen in total CD4+ T cells using a library of open chromatin obtained from the same cells as input. This cell pool is comprised of diverse CD4+ T cell subtypes, many of which are likely to utilize unique CRE subsets. Though it is possible to glean subtype-specific CRE function based on accessibility and chromatin modifications, we cannot rule out that the STARR-Seq functional CRE identified herein are functional in only some CD4+ subtypes. This limitation is an ongoing technical challenge for screening methods, including STARR-Sequencing, as it requires high read-density coverage and tens of millions of cells per biologic replicate. With the advent of single-cell sequencing and single-cell CRISPR screens, these limitations may become less technically and fiscally burdensome.

Broadly, two mechanisms of silencer action have been proposed: 1) competing DNA binding between activating and repressing proteins and 2) recruitment of negative regulators, such as HDACs, with subsequent formation of heterochromatin domains.(66, 67) Although we do observe local heterochromatin mark enrichment at functional NREs, use of an accessible chromatin library as input precludes analysis of functional NREs within expansive heterochromatin marked domains. Rather, we demonstrate that functional NREs can exist in open chromatin and even within ‘activated’ chromatin environments. These observations suggest some competition between transcription factors with opposing functions. The presence of PolII at NREs, but lower presence of activated S5-PolII at NREs compared to enhancers, suggests that regulation of polymerase may be a silencing mechanism used by at least some NREs. Though we did not identify a central unifying repressor in CD4+ T cells, we found that these NREs were likely to recruit transcription factors and co-activators with known repressor and nucleosome-repositioning function. Other silencer screens corroborate this widely varied transcription factor and motif utilization across the functional silencer elements; no unifying chromatin marks or transcription factor binding profile of functional NREs has been identified to date.(1–3, 45, 67)

This screen demonstrates striking differences in patterns of epigenetic enrichment between enhancers and NREs. Other screens have identified functional silencer activity to be associated with H3K36me3 and H3K9ac, marks of active transcription and enhancers. We also found enrichment of these marks at NREs (**S1**); analysis of functional silencers in K562 cells by Pang *et al*. 2020 also found that silencers were marked with H3K27ac and alternate histone H2A.Z. Intriguingly, some STARR-Seq NREs existed within active chromatin environments, including super-enhancers (**S4)**. Previously, a detailed analysis of one super-enhancer found that some of its constituent elements function as silencers.(68) These multimodal ATAC-Seq sites may be multifunctional, providing both positive and negative regulatory information to nearby genes in different cellular contexts. Further study is required to refine the *in situ* function of these complex enhancer:NRE relationships at the level of individual genes.

Our approach queried the function of accessible CREs for enhancer and negative regulatory activity from a library of open chromatin; this assay excluded interrogation of CRE within heterochromatin and inaccessible chromatin environments. Screens for silencers from libraries of repressed chromatin, performed in Drosophila and other Hi-C experiments, suggest that at least some CREs within H3K27me3 domains are also functional as NREs.(41, 45) Intriguingly, we also found some heterochromatin mark enrichment at functional NREs in CD4+ T cells, despite a paucity of this mark at sites of open chromatin in mature CD4+ T cells. We demonstrate that NREs and enhancers display key differences in local chromatin accessibility. STARR-Seq functional NREs possess centrally positioned nucleosome, which may drive the strong central enrichment that we observed for the histone marks H3K27ac and H3K4me3 (**S5**). It is known that nucleosome repositioning by transcription factors can be repressive when placed in unfavorable locations for transcription.(69) A direct investigation of silencers that form heterochromatin found that silencing requires precise nucleosome positioning and can be sensitive to nucleosome repositioning.(70) Other investigations find that nucleosome positioning is important for silencing of heterochromatically marked genes more broadly.(71, 72) As our assay tests putative CREs outside of their native chromatin environment, some NREs may contain intrinsic nucleosome-positioning information, or be permissive to nucleosome occupancy. These possibilities could support a potential silencing mechanism related to local chromatin accessibility changes that prevent transcriptional elongation. Alternatively, STARR-Seq may uncover functional silencers that are endogenously occupied by nucleosomes. Intriguingly, we demonstrate that these NREs are functional from numerous genomic locations: intronically (STARR-Seq), upstream of a promoter (Luciferase), and in long-range chromatin looping *in situ* (**3E**). Together this suggests that their mechanism is not limited to unfavorable nucleosome placement blocking intron transcription.

Others have speculated that NREs may have alternate enhancer functions in unrelated cell types. We also demonstrate that CD4+ functional NREs resemble the enhancer phenotype in other cells due to their enrichment in FANTOM5, and in the enhancer histone marks H3K4me1 and H3K27ac. Though functional evidence is needed to support this hypothesis, we posit on the basis of our own findings that NRE function may also be incorrectly assumed as enhancing. Although it is tempting to ascribe enhancer activity to histone mark enrichment alone, we and others demonstrate that functional NREs exist within chromatin environments traditionally associated with enhancers. This challenge extends to many enhancer and silencer imputation attempts, which mainly leverage chromatin state. Indeed, we found that imputed silencer location in CD4+ T cells from SilencerDB had poor enrichment of Lenti-STARR NREs and enhancers (**S6**).(73) We emphasize the need for further investigation into NRE behavior across developmental time periods and transitions and specifically into whether NREs are repurposed as enhancers or simply maintain repressive function.

## Acknowledgements

We thank Benjamin Wronowski for Tn5 preparation. We also thank Matt Weirauch for his assistance with RELI.

## Description of Supplemental Files

Supplemental File 1: Supplemental Figures and Figure Legends

Figure S1: ChIP-Seq of Histone Marks Previously Reported to be enriched at silencers

Figure S2: Transcription Factor ChIP-Seqs Performed in CD4+ T Cells

Figure S3: Expanded Motif Enrichment at Cis-Regulatory Elements

Figure S4: Cis-Regulatory Enrichment Within SuperEnhancers

Figure S5: Cis-Regulatory Element Histone ChIP-Seq Enrichment with Nucleosome Positive and Negative Clustering

Figure S6: Silencer Database (SilencerDB) Intersection of Cis-Regulatory Elements

Figure S7: Expression of Genes Nearest to Cis-Regulatory Elements

Supplemental Table 1: Primers and dsDNA sequences used for luciferase validation Supplemental Table 2: Luciferase Values for NREs and enhancer single-site validation. Supplemental Table 3: CRE locations described by Lenti-STARR-Seq

1. MACS2 narrowpeak output for all enhancer sites
2. Final significant enhancers
3. FAST-NR output for all NRE sites
4. Final significant NREs
5. MACS2 narrowpeak output for all input plasmid peaks

Supplemental Table 4: Utilized datasets

