## Supplemental Information for "Cis-Regulatory Atlas in Primary Human CD4+ T Cells"

**Supplemental Materials and Figure Legends**

Supplemental File 1: Supplemental Figures and Figure Legends
- Figure S1: ChIP-Seq of Histone Marks Previously Reported to be enriched at silencers
- Figure S2: Transcription Factor ChIP-Seqs Performed in CD4+ T Cells
- Figure S3: Expanded Motif Enrichment at Cis-Regulatory Elements
- Figure S4: Cis-Regulatory Enrichment Within SuperEnhancers
- Figure S5: Cis-Regulatory Element Histone ChIP-Seq Enrichment with Nucleosome Positive and Negative Clustering
- Figure S6: Silencer Database (SilencerDB) Intersection of Cis-Regulatory Elements
- Figure S7: Expression of Genes Nearest to Cis-Regulatory Elements

Supplemental Table 1: Primers and dsDNA sequences used for luciferase validation

Supplemental Table 2: Luciferase Values for NREs and enhancer single-site validation.

Supplemental Table 3: CRE locations described by Lenti-STARR-Seq

1. MACS2 narrowpeak output for all enhancer sites
2. Final significant enhancers
3. FAST-NR output for all NRE sites
4. Final significant NREs
5. MACS2 narrowpeak output for all input plasmid peaks

Supplemental Table 4: Utilized datasets

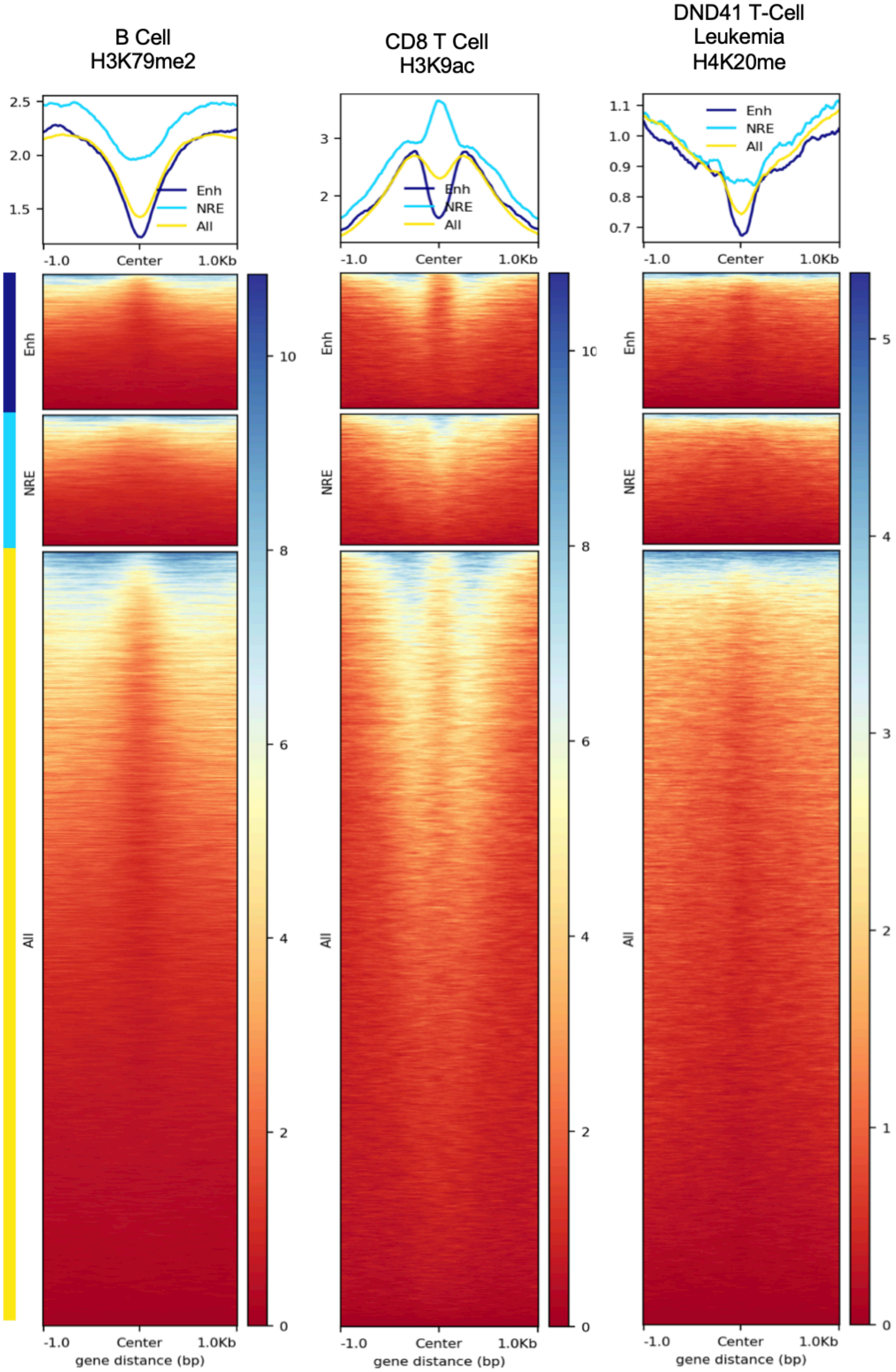


**Supplemental Figure 1: ChIP-Seq of Previously Proposed Identified Silencer Enriched Histone Marks.** Enrichment of transcription factors or histone ChIP-Seq experiments in non-CD4+ T cells. The selected histones demonstrate enrichment at negative regulatory elements (NREs) in other silencer studies (Pang *et al.* 2020) (Gisselbrecht *et al.* 2020).

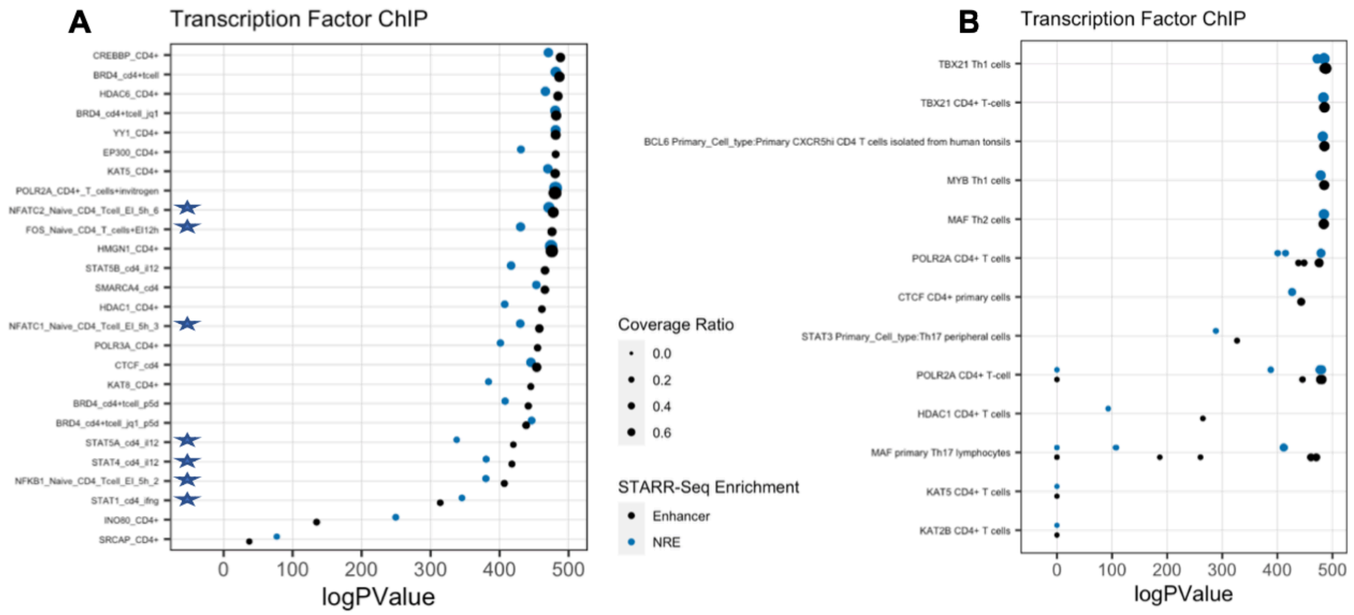


**Supplemental Figure 2: Transcription Factor ChIP-Seq Performed in CD4+ T Cells.** RELI enrichment LogP-value of ChIP-Seq experiments performed in human CD4+ T cells across STARR-Seq–identified Cis-regulatory elements (CREs). Star indicates the experiment was performed in activated cells. **A)** Datasets included in the original RELI manuscript and **B)** Datasets from GEO database.(48)

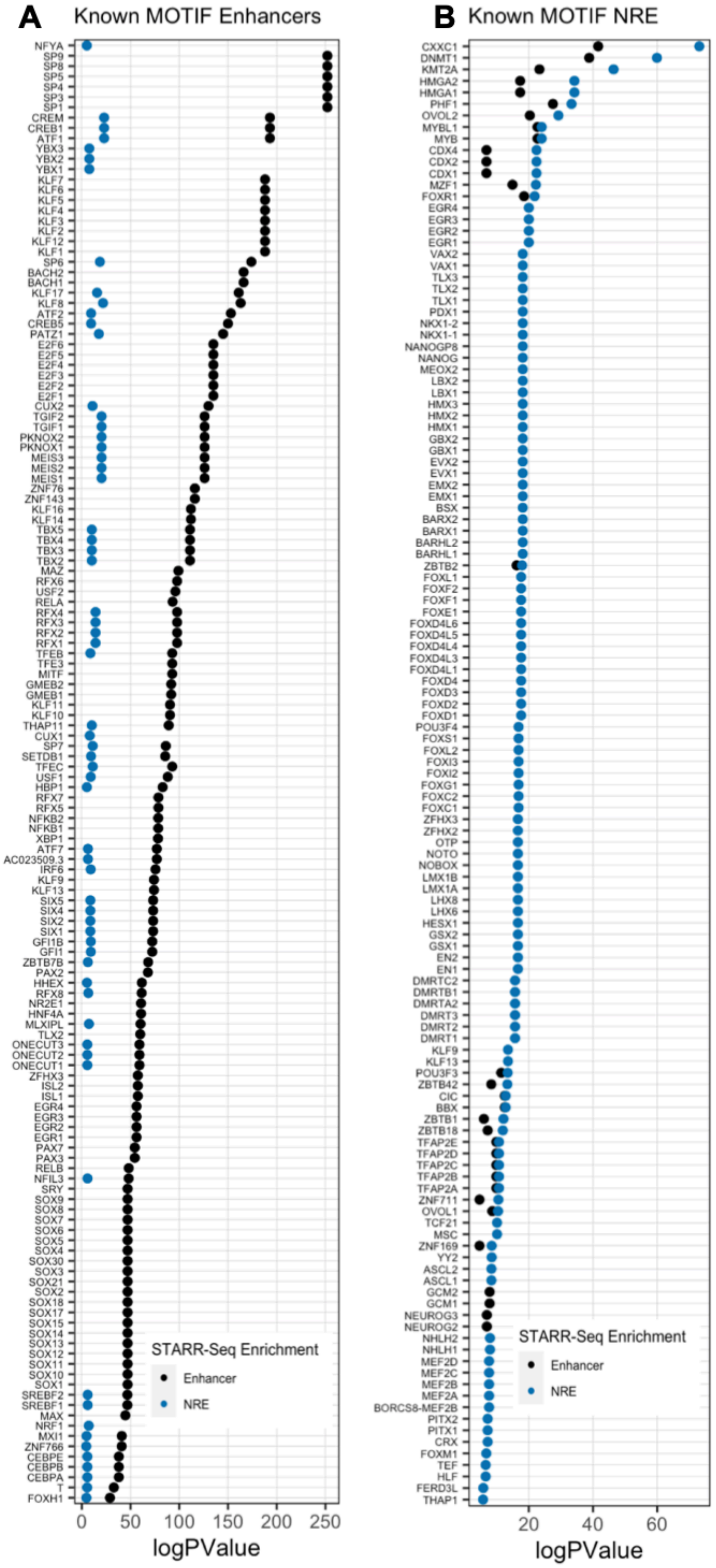


**Supplemental Figure 3: Expanded Motif Enrichment at Cis-Regulatory Elements.** Motif enrichment greater across **A)** Enhancers and **B)** negative regulatory elements (NRE)s at known motifs (n=7704) from Cis-BP Database known Motif database.

A

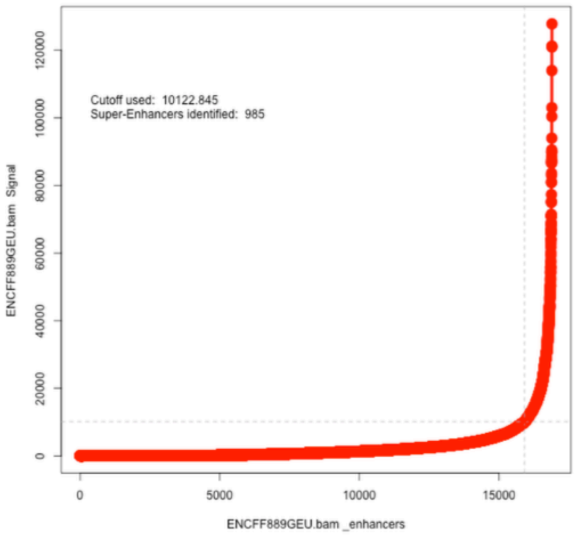


B

| Chi-Squared p-value | ATAC-Seq All Peaks |
|---|---|
| Enh | 6.353e-13 |
| NRE | <2.2e-16 |

**Supplemental Figure 4: Cis-Regulatory Enrichment Within SuperEnhancers A)** SuperEnhancer ranks are called using ROSE algorithm with H3K27ac ChIP-Seq experiment ENCFF889GEU. **B)** 2x2 Chi-squared test for the difference in the number of overlaps of SuperEnhancers with Enhancer [or negative regulatory elements (NREs)] versus all ATAC-Seq input peaks.

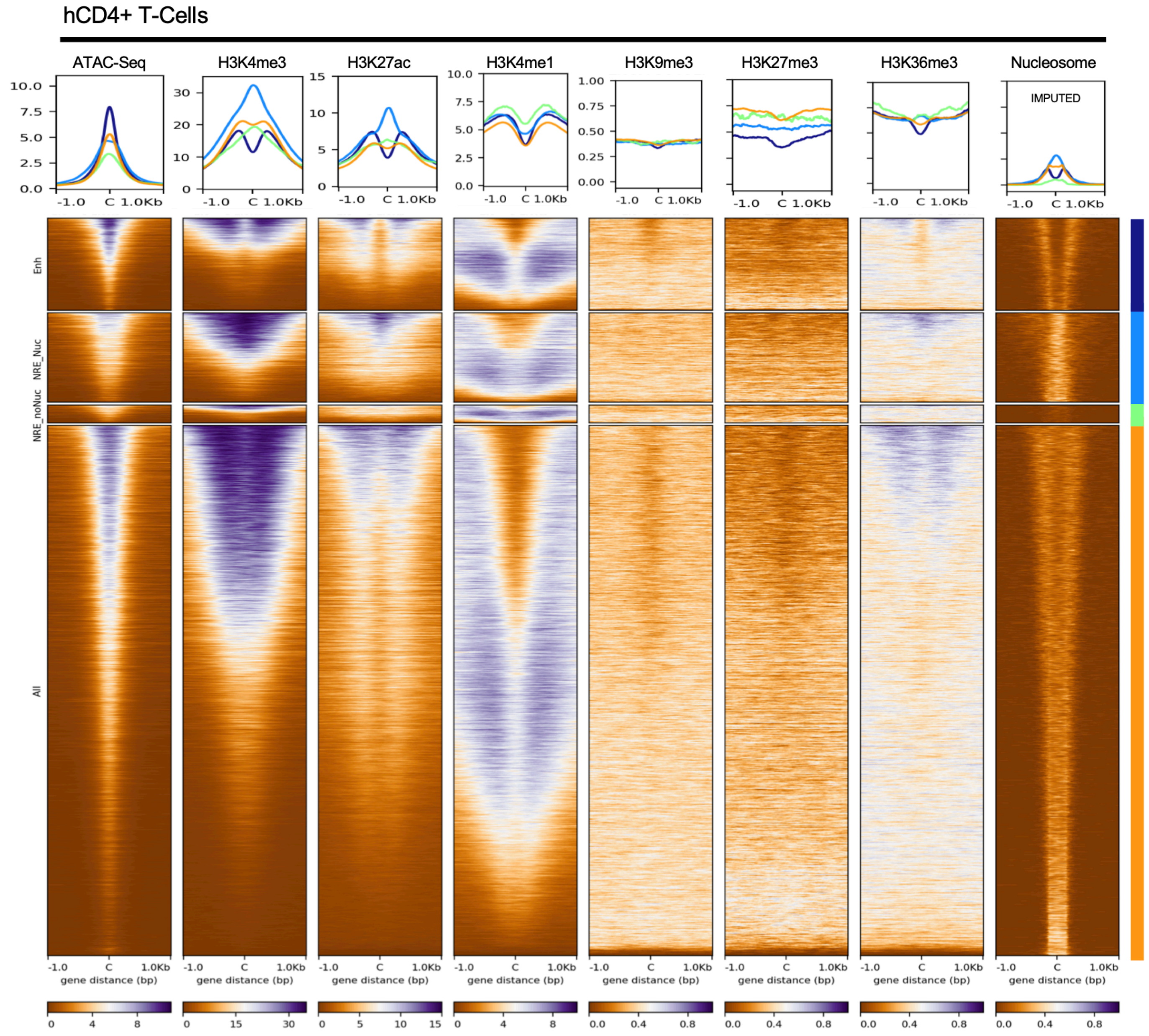


**Supplemental Figure 5: Cis-Regulatory Element Histone ChIP-Seq Enrichment with Nucleosome-Positive and Nucleosome-Negative Clustering.** Tag density plots of mean intensity and heatmaps of ChIP-Seq or ATAC-Seq performed in human CD4+ T cells plotted against STARR-Seq–identified Enhancers, negative regulatory elements (NREs), and 'All' open peaks from input. (ENCODE). Nucleosome location in resting CD4+ T cells imputed from NucleoATAC. Tag density of mean signal intensity displayed for histone enrichment and of imputed nucleosome position. NRE sites are k-means (n=2) clustered by imputed nucleosome signal.

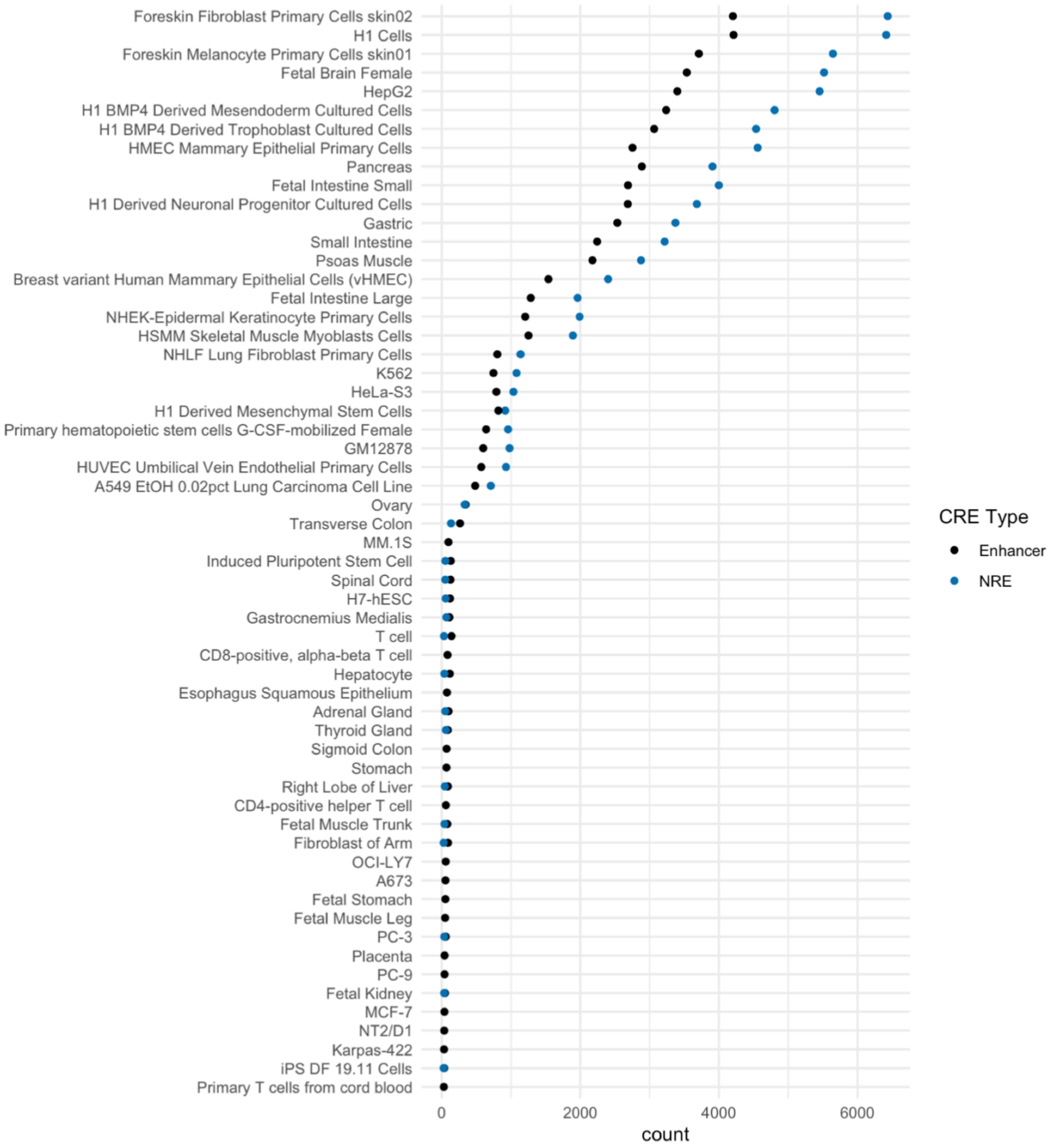


**Supplemental Figure 6: Silencer Database (SilencerDB) Intersection of Cis-Regulatory Elements.** The number of Lenti-STARR-Seq–identified Cis-regulatory elements (CREs) intersected with predicted silencers in SilencerDB.(73)

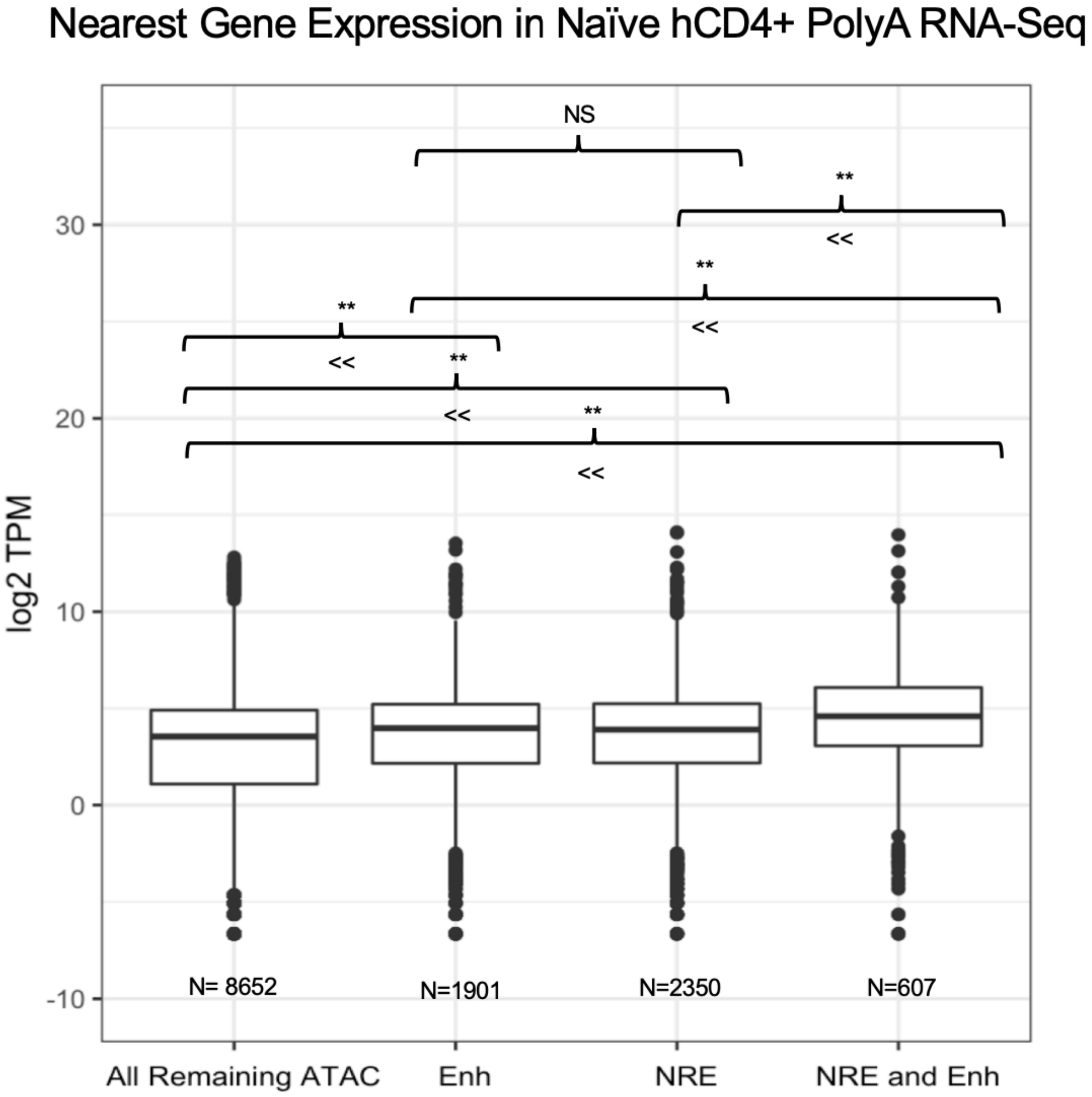


**Supplemental Figure 7: Expression of Nearest Genes to Cis-Regulatory Elements.** Transcripts per million (TPM) in human (h)CD4+ PolyA RNA-Seq (ENCSR545MEZ) at TSS within 10 kb of genes with either: both Enhancer and negative regulatory elements (NREs) near, Enhancer only, NRES only, or 'All Other' open peaks from input. Boxplot displays median and interquartile range. (** P-value < 0.01 by non-parametric Kruskal-Wallis two-sided test with Holm adjustment for multiple comparisons)
